# Mutually exclusive teams-like patterns of gene regulation characterize phenotypic heterogeneity along the noradrenergic-mesenchymal axis in neuroblastoma

**DOI:** 10.1101/2023.09.03.556132

**Authors:** Manas Sehgal, Sonali Priyadarshini Nayak, Sarthak Sahoo, Jason A Somarelli, Mohit Kumar Jolly

**Affiliations:** Centre for BioSystems Science and Engineering, Indian Institute of Science, Bangalore, 560012, India; Max Planck School Matter to Life, University of Göttingen, Friedrich-Hund-Platz 1, 37077 Göttingen, Germany; Department of Medicine, Duke University, Durham, NC 27708, USA

**Keywords:** Phenotypic heterogeneity, Noradrenergic-Mesenchymal Transition, Mesenchymal-Noradrenergic Transition, GD2, Neuroblastoma

## Abstract

Neuroblastoma is the most frequent extracranial pediatric tumor and leads to 15% of all cancer-related deaths in children. Tumor relapse and therapy resistance in neuroblastoma are driven by phenotypic plasticity and heterogeneity between noradrenergic (NOR) and mesenchymal (MES) cell states. Despite the importance of this phenotypic plasticity, however, the dynamics and molecular patterns associated with these bidirectional cell-state transitions remain relatively poorly understood. Here, we analyze multiple RNA-seq datasets at both bulk and single-cell resolution, to understand the association between NOR- and MES-specific factors. We observed that NOR-specific and MES-specific expression patterns are largely mutually exclusive, exhibiting a “teams-like” behavior among the genes involved, reminiscent of our earlier observations in lung cancer and melanoma. This antagonism was noticed also in the association of NOR and MES phenotypes with metabolic reprogramming and with immunotherapy targets PD-L1 and GD2, and in experimental perturbations driving the NOR-MES and/or MES-NOR transition. Further, this “teams-like” patterns was seen only among the NOR- and MES-specific genes, but not in housekeeping genes, possibly highlighting a hallmark of network topology enabling cancer cell plasticity.

## Introduction

Neuroblastoma (NB) is the most frequent pediatric solid tumor that represents 6%-10% of all childhood tumors and accounts for approximately 15% of all cancer deaths in children. It almost exclusively occurs in early childhood; the median age for its diagnosis is about 18 months, and approximately 40% of NB patients are younger than 1 year at diagnosis. It is a neural crest-derived malignancy that manifest along the sympathetic nervous system (Brodeur, 2003; Johnsen et al., 2019; Park et al., 2010). NB is marked by considerable clinical variability, with the disease spectrum varying from spontaneous regression without needing any treatment to a treatment-resistant tumor exhibiting metastatic dissemination (Johnsen et al., 2019). Treatment usually includes high-dose chemotherapy, surgical resection, radiation therapy and immunotherapy (anti-GD2 monoclonal antibodies), with most targeted therapy-based inhibitors still in clinical trials (Greengard, 2018; Zage, 2018). Despite intensive therapy, NB patients have high mortality rates, and relapsed patients frequently develop treatment resistance. The median survival for high-risk relapsed neuroblastoma is only 11 months, and there are no curative options for relapsed patients, underscoring the urgent need to improve current treatment regimens (Shendy et al., 2022).

NB exhibits extensive transcriptional heterogeneity, akin to reports in other cancer types (Pillai and Jolly, 2021; Sharma et al., 2019; Su et al., 2019). A low mutational burden in the typical NB genome highlights the role of non-genetic heterogeneity in enabling therapy resistance (Shendy et al., 2022). Over five decades ago, neuroblastoma cells were shown to exhibit two distinct phenotypes *in vitro*: the N-type (neuroblast) and S-type (substrate-adherent). The N-type grew as focal aggregates with short neurotic processes and attached poorly to substrate, while S-type were largely flattened, attached strongly to substrate, and resembled non-neuronal precursors. Interestingly, both these cell types could interconvert spontaneously and bidirectionally (Gautier et al., 2021), reminiscent of phenotypic plasticity reported in other cancers (Bhatia et al., 2019; Chauhan et al., 2021). Later, a morphological and biochemical intermediate (I-type cells) were proposed; they had the highest tumor-forming abilities, expressed stem cell markers (Gautier et al., 2021; Walton et al., 2004), and were postulated to be a possible precursor to both N-type and S-type cells. Extensive analysis of developmental origins of neuroblastoma have drawn parallels between sympathoblasts – the bipotent cells that can generate both mesenchymal and neuronal phenotypes – and I-type cells. During development, sympathoblasts can give rise to both neuronal (post-ganglionic sympathetic neurons, chromaffin cells) and mesenchymal (mesenchymal stem cells, fibroblasts, Schwann cells, fibroblasts) cell types (**Figure 1**) (Zeineldin et al., 2022). Further molecular characterization led to N-type cells being referred to as noradrenergic (NOR) phenotype, while S-type being renamed as mesenchymal (MES) phenotype (Gautier et al., 2021).

**Figure 1.**
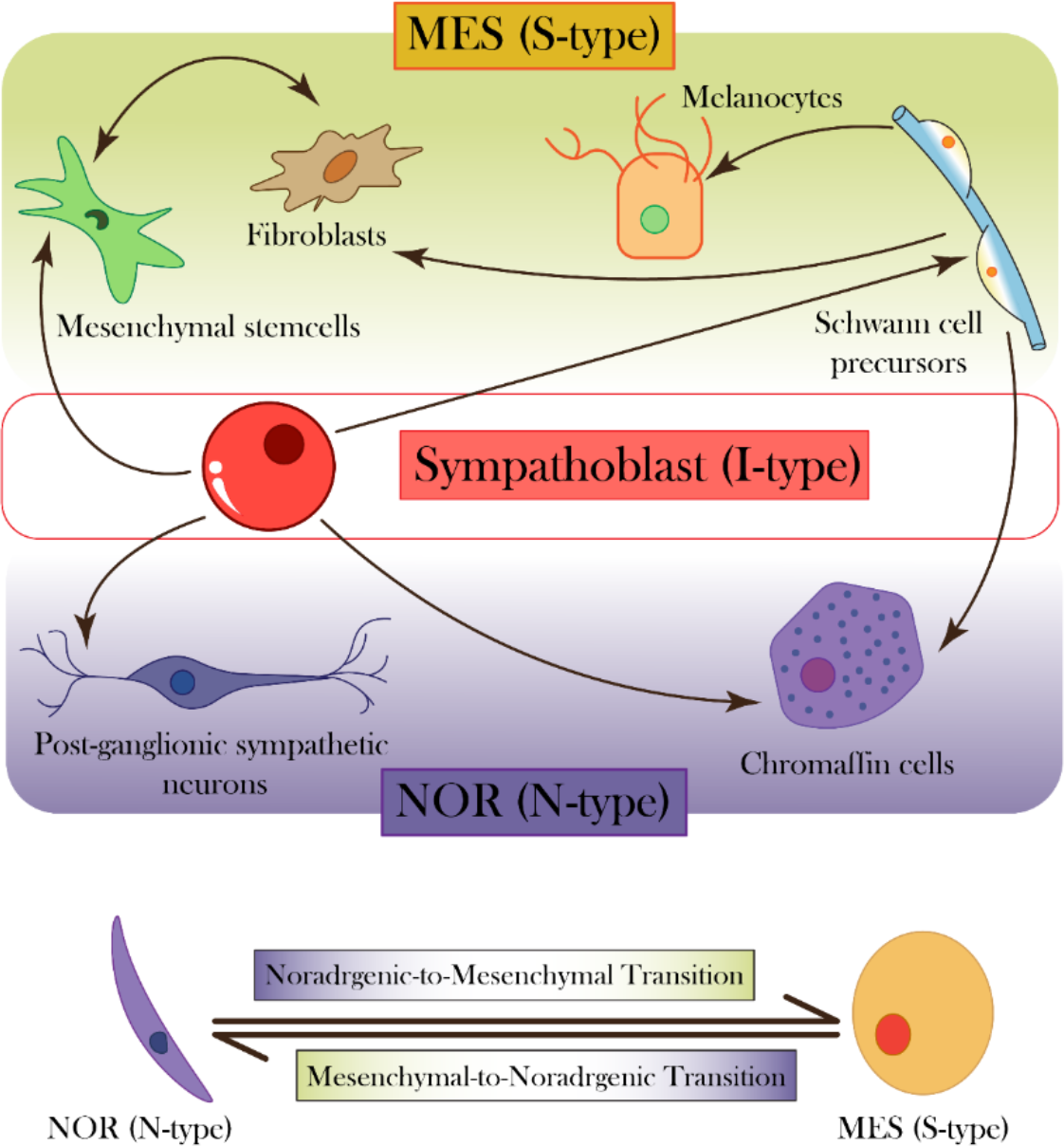
Neuroblastoma phenotypes and corresponding developmental states. (Top) Sympathoblast (I-type) serves as a progenitor of many cell types shown in NOR (N-type) and MES (S-type) lineage, at different stages of lineage differentiation. (Bottom) NOR and MES cells can switch back and forth, indicating reversible cell-state transitions.

Compared to NOR cells, MES cells are resistant to standard chemotherapeutic agents used for neuroblastoma patients – cisplatin, doxorubicin, and etoposide, and are enriched upon treatment of the heterogeneous cell line, SK-N-SH (containing both NOR and MES subpopulations) with these drugs (Gautier et al., 2021). Consistently, they are also enriched in patients with relapse, as identified by single-cell analysis of primary and relapsed tumors (Shendy et al., 2022). This enrichment can be enabled by selection of pre-existing MES cells and/or induction of NOR to MES cell-state transition. Single-cell subclones established from NB cell lines could repopulate both cell-states *in vitro*, indicating reversible dynamic transitions. Such spontaneous stochastic bidirectional interconversion was also seen *in vivo* (Lecca et al., 2018; Thirant et al., 2023; Van Groningen et al., 2017). Therefore, analogous to the well-studied phenomenon of epithelial-mesenchymal transition (EMT) and its reverse mesenchymal-epithelial transition (MET) (Kahlert et al., 2017; Lourenco et al., 2020; Pastushenko and Blanpain, 2019), NB displays phenotypic transitions between the NOR and MES phenotypes - noradrenergic-mesenchymal transition (NMT) and mesenchymal-noradrenergic transition (MNT) (**Figure 1**).

NMT/MNT can drive extensive phenotypic plasticity and heterogeneity reported in NB, highlighting the importance of non-genetic mechanisms, such as epigenetic and transcriptomic reprogramming in enabling disease progression and relapse (Boeva et al., 2017; Thirant et al., 2023; Van Groningen et al., 2017). While the epithelial-hybrid-mesenchymal spectrum has been thoroughly investigated with respect to its dynamics and impact on hallmarks of cancer (Pastushenko and Blanpain, 2019; Sahoo et al., 2021; Simeonov et al., 2021; Tripathi et al., 2020), our understanding of NMT/MNT is relatively limited, and additional research is needed to uncover the underlying patterns of phenotypic plasticity and heterogeneity in NB cell populations. A deeper understanding of the dynamics of cell fate switching in NB could help pinpoint new biomarkers or treatments to target aggressive cell states in NB.

Here, we found a mutually exclusive expression pattern between two sets of TFs in NB, one driving a NOR phenotype and the other enabling a MES one, thus indicating a teams-like behavior between gene expression patterns that determine these phenotypic cell states. This ‘teams’ behavior can underlie the phenotypic plasticity, and is largely unique to gene lists associated with noradrenergic/mesenchymal axis in NB. We also demonstrated that the NOR- and MES-associated gene lists could reproduce cell-state transitions in NB. Further, our meta-analysis of NB datasets confirmed an antagonistic enrichment of the two phenotypes in bulk transcriptomic datasets along with the enrichment of a PD-L1, glycolytic, and fatty acid oxidation (FAO) programs with a mesenchymal state of NB cells. Further, we found GD2, a cell surface marker abundantly expressed in NB tumors, is associated with a NOR phenotype both at bulk and single cell levels. Our analysis elucidates functional and molecular differences between NOR and MES phenotypes prevalent in NB.

## Materials and Methods

### Software, and data and code availability

All computational and statistical analyses have been performed using R (version 4.2.1) and Python (version 3.10.2). Codes used are available at https://github.com/Manas-Sehgal/NB_heterogeneity

### Transcriptomic data retrieval and pre-processing

Publicly accessible bulk and single-cell RNA-sequencing datasets were downloaded from NCBI-GEO) repository. Raw gene count matrices were normalized for gene length and transformed to Log2(TPM) (transcripts-per-million) values. Single-cell RNA-sequencing gene-length normalized data (scRNA-seq) was imputed to reduce sparsity using ‘Rmagic’ package (van Dijk et al., 2018). Microarray datasets were downloaded from GEO using the GEO*query* R Bioconductor package. A total of 75 NB datasets were used for meta-analysis (**Table S3)**. Probe-wise gene expression matrices and their corresponding platform annotation files were used to map the probes to each gene. If two probes matched to the same gene, their mean expression values were used. The resultant gene-wise expression matrices were then log2 transformed.

### Principal component analysis and K-means clustering

Principal component analysis (PCA) was used for dimensionality reduction of simulation and transcriptomic data using ‘Scikit-learn’ python library. K-means clustering was performed on samples in each bulk dataset using the ‘Scikit-learn’ python library to create two groups (K = 2) based on their segregation along principal components.

For gene swapping analysis, PCA was performed on gene-expression matrices before and after swapping genes from the wild-type 26 NOR/MES gene list was done using ‘prcomp’ function in R. List of housekeeping genes for this analysis was obtained from a previous study (**Table S2**) (Eisenberg and Levanon, 2013).

### Gene signatures and gene set enrichment analysis

A total of 14 NOR-specific and 12 MES-specific genes (**Table S1**), and extended lists of 369 NOR-specific and 485 MES-specific genes from a previous study were used (Van Groningen et al., 2017). Gene set enrichment analysis (GSEA) was performed on K-means-generated clusters to examine the enrichment of the noradrenergic and mesenchymal genesets using GSEAPY python library (Subramanian et al., 2005a). Gene signatures for Hallmark EMT, fatty acid oxidation (FAO) and glycolysis pathways, Hallmark G2M checkpoint geneset, WP cell cycle, Biocarta cell cycle and KEGG cell cycle were obtained from molecular signatures database (MSigDB) (Liberzon et al., 2011). Single-sample gene set enrichment analysis (ssGSEA) was performed to obtain sample-wise normalized enrichment scores (NES) for each pathway for bulk transcriptomic data and AUCell was performed on the imputed scRNA-seq gene matrices (Aibar et al., 2017). Partial EMT and Programmed death-ligand 1 (PD-L1) scores were obtained using previous gene lists (Puram et al., 2017; Sahoo et al., 2021). All gene lists used in this study are included in **Table S2**.

### Gene correlation matrices and J-metric calculation

Spearman’s correlation coefficients were used to assess the pairwise association between MES-specific and NOR-specific genes. R package ‘ggcorrplot’ was used to plot the correlation matrices. Normalized expression values of genes corresponding to NOR and MES phenotypes (shown in Figure 1) were used to calculate J-metric value for each dataset. The sum of Spearman correlation coefficients between genes of different teams were subtracted from the sum of correlation coefficients of genes belonging to the same team. Only significant correlations (p < 0.05) were considered for this calculation.

## Results

### NB phenotypic heterogeneity includes the existence of two distinct phenotypes and their corresponding ‘teams’ of genes

To characterize phenotypic heterogeneity in NB, we first collated a set of 14 NOR-specific and 12 MES-specific genes that regulate and/or correspond to noradrenergic and mesenchymal cell-states (**Table S1**) from a previous study (Van Groningen et al., 2017). In our prior studies of epithelial-mesenchymal plasticity in cancer, we have observed that sub-networks of specific genes exist as coordinated “teams” to regulate the dynamics of cellular plasticity (Hari et al., 2022). Players within a “team” activated each other, while those across “teams” inhibited each other directly or indirectly, thus the presence of “teams” can lead to mutually exclusive expression patterns in transcriptomic data. For example, we have pinpointed the epithelial “team” comprised of players such as microRNA-200, OVOL2, GRHL2, and the mesenchymal “team” comprised of players such as ZEB1, SNAIL1 and TWIST (Chakraborty et al., 2021). Similar “teams” behaviour was seen in both the underlying network structure as well as in transcriptomic datasets in additional cancer types, where two mutually antagonistic “teams” promoted invasive/proliferative heterogeneity in melanoma and neuroendocrine/non-neuroendocrine heterogeneity in small cell lung cancer (Chauhan et al., 2021; Pillai and Jolly, 2021). With these features in mind, we sought to understand if the genes involved in regulating the NOR and MES phenotypes in NB were organized in “teams” as well.

To do this, we then computed the pairwise correlations between these genes in multiple bulk transcriptomic datasets (denoted by GSE IDs) comprising of NB cell lines and primary tumor samples – GSE9169 (n=86) (Nishida et al., 2008), GSE17714 (n=22) (Fardin et al., 2010), GSE28019 (n=24), GSE64000 (n=8) (Middelbeek et al., 2015), GSE66586 (n=10) (Gu et al., 2015), GSE78061 (n=29), GSE19274 (n=138) (Cole et al., 2011), GSE44537 (n=6) (Ikegaki et al., 2013) and GSE73292 (n=6) (Debruyne et al., 2016). We observed that expression levels of 14 NOR-specific genes were mostly positively correlated with each other, but negatively correlated 12 MES-specific genes. Similarly, the 12 MES-specific genes correlated positively with each other, but negatively with 14 NOR-specific ones (**Figure 2**). This consistent pattern across datasets indicates the existence of two “teams” of players wherein one set of genes corresponds to a NOR phenotype and the other set corresponds to the MES phenotype. Thus, such mutual exclusivity between the two “teams” characterizes NB heterogeneity patterns.

**Figure 2.**
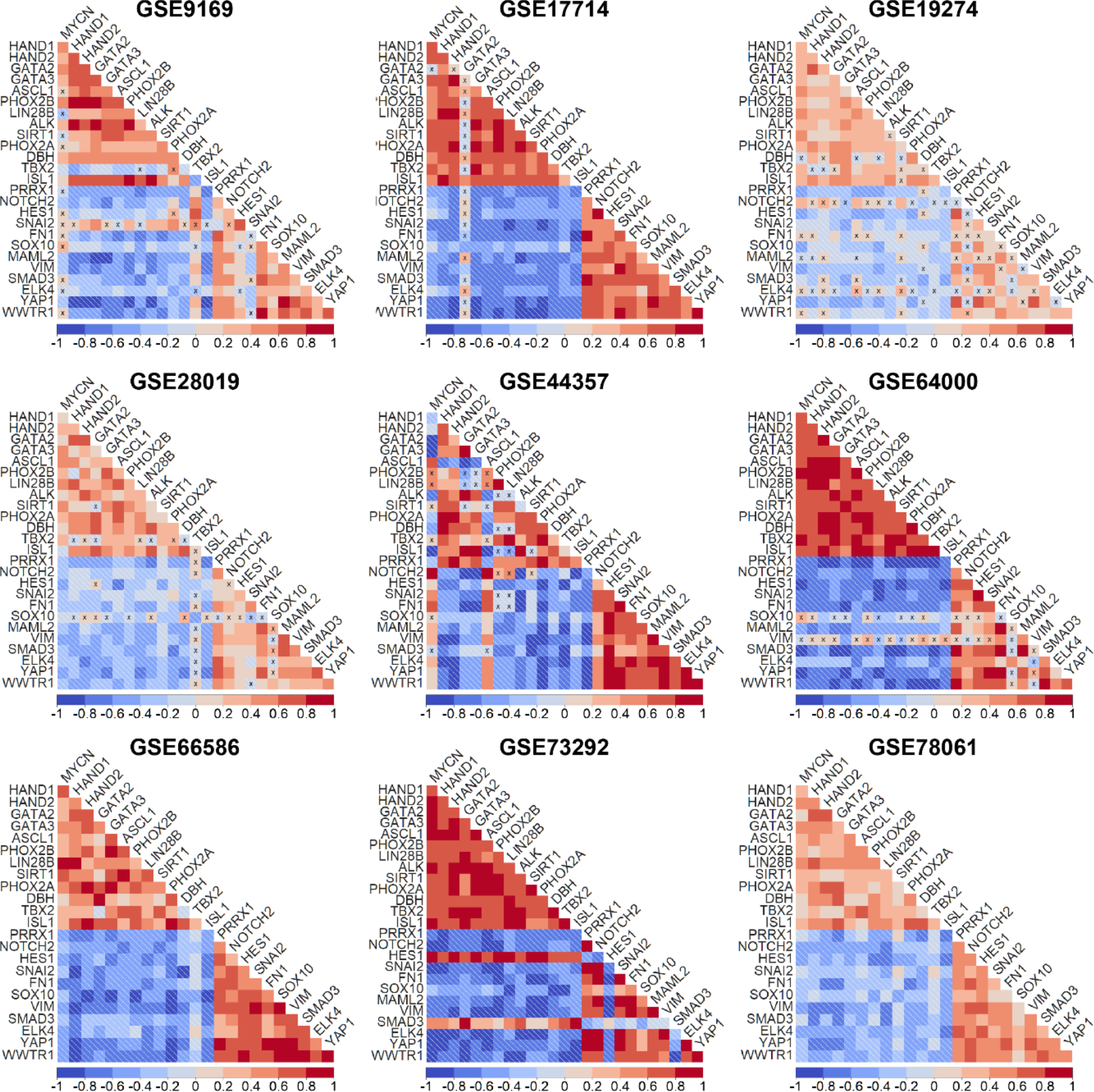
Prevalence of ‘team-like’ behavior corresponding to the two major phenotypes — noradrenergic and mesenchymal. Correlation matrices illustrating pairwise Spearman’s correlation of NOR-specific and MES-specific genes for GSE9169 (n = 86), GSE17714 (n = 22), GSE28019 (n =24), GSE64000 (n = 8), GSE66586 (n = 10), GSE78061 (n = 29), GSE19274 (n = 138), GSE44537 (n = 6) and GSE73292 (n = 6). All pairwise correlations are significant (p< 0.05) except the ones indicated by ‘X’. The color bar legend depicts the values of the correlation coefficient. ‘n’ denotes the number of samples in each dataset.

To better understand this “teams” behavior, we applied dimensionality reduction to the transcriptomic datasets. In our previous analysis of cell-fate decision networks of EMT and pluripotency, we observed that the underlying “teams” structure in a network can reduce the dimensionality of the system and could segregate the phenotypes distinctly along principal component 1 (PC1) in a principal component analysis (PCA) (Hari et al., 2023). When projected on their first two principal components (PCs), we observed the segregation of samples into two distinct clusters along PC1 across the datasets shown, further highlighting the distinct, teams-like behavior of these regulatory pathways (**Figure 3A-E**). Next, we applied K-means clustering (K=2) to samples in each dataset to identify two clusters (represented by yellow and purple color) and calculated the enrichment of the reported 485-gene MES and 359-gene NOR signatures in these two segregated clusters using GSEA (Subramanian et al., 2005b). We noticed a striking pattern across the datasets, albeit to varying extents, that while one of the two clusters showed enrichment of NOR gene list, the other cluster showed enrichment of the MES gene set (**Figure 3A-E, S1**). Also, the extent of enrichment of NOR and MES phenotypes was anti-correlated across datasets (**Figure 3F, Table S4**). Together, these observations indicate that the two observed clusters in each dataset can be mapped onto the NOR and MES phenotypes.

**Figure 3.**
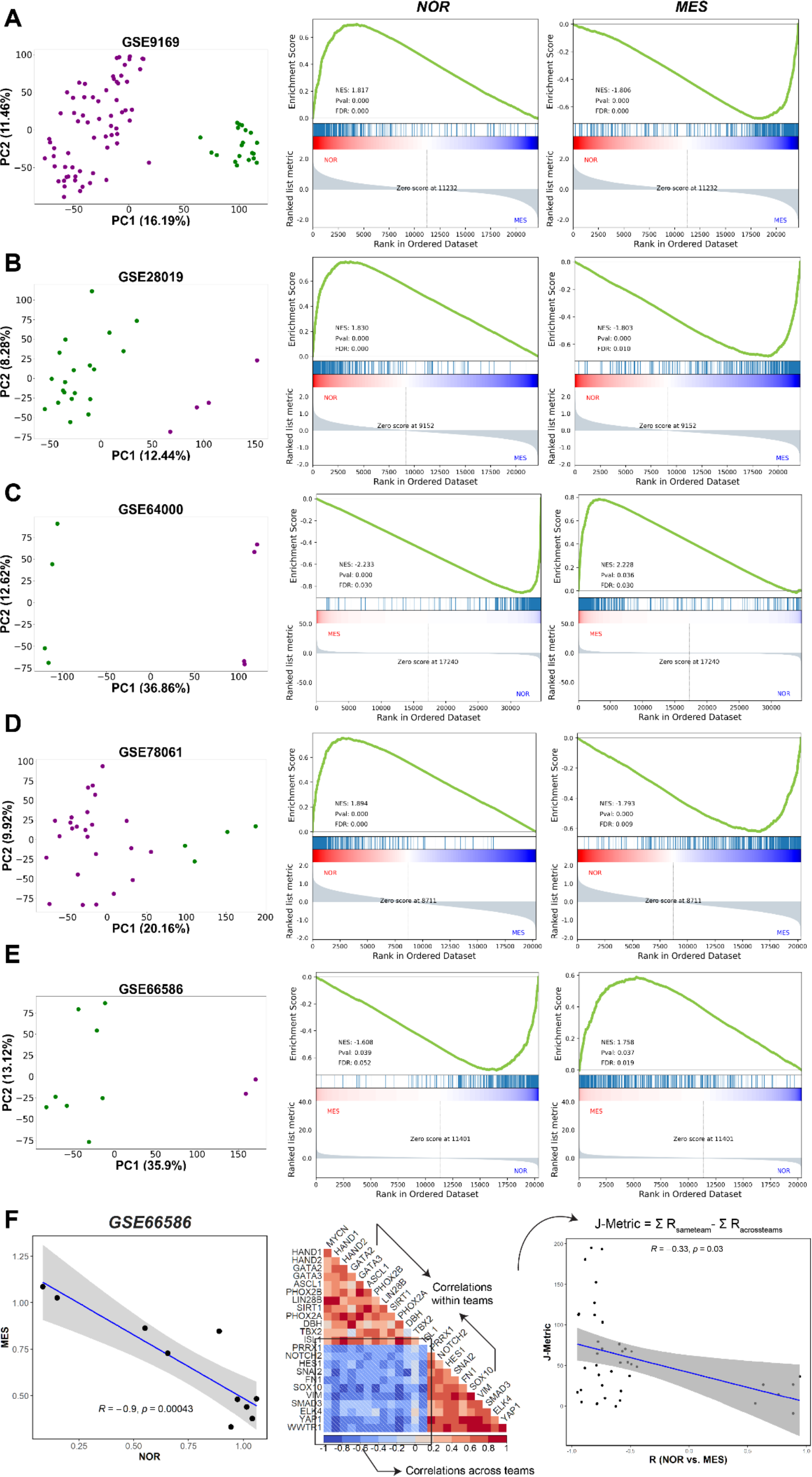
Two distinct phenotypes—mesenchymal and adrenergic – are observed in NB. **A)** PCA plot showing two distinct classes of samples exist in multiple datasets. K-means clustering for K = 2 yields two clusters (shown in green and violet color) along principal component 1 (PC1). GSEA for the mesenchymal (MES) geneset (middle) and adrenergic (NOR) geneset (right) confirms that these two clusters correspond to respective phenotypes for **A)** GSE9169 (n = 86), **B)** GSE28019 (n =24), **C)** GSE64000 (n = 8), **D)** GSE66586 (n = 10) and **E)** GSE78061 (n = 29). ‘n’ stands for number of samples in each dataset. Percentage variance explained by each PC is indicated along the respective axes. **F)**. Scatter plot depicting NOR (x-axis) and MES (y-axis) ssGSEA scores of samples of GSE66586 (left). Correlation matrix for GSE66586 representing the calculation of J-metric using the correlations between genes shown in Figure 1 (middle) and scatter plot depicting the Spearman correlation coefficient ‘R’ between NOR and MES scores across 75 bulk datasets (x-axis) and corresponding J-metric (y-axis). Only 42 datasets in which NOR and MES show significant correlation (p <0.05) have been included (**Table S6**).

Further, we evaluated whether the segregation of samples based on a relatively small gene list (14 NOR-specific and 12 MES-specific) correlated with the clustering obtained through a much larger dimensionality of the system investigated (359-gene NOR signature, 485-gene MES signature). To this end, we used a set of 75 transcriptomic NB datasets for this analysis (**Table S3**). Previously, we defined a J-metric to quantify the extent of “teams” behavior in each transcriptomic dataset by analyzing the values of a corresponding pairwise correlation matrix (Chauhan et al., 2021). The larger the J-metric, the stronger the “teams” behavior in terms of mutual exclusivity and antagonism between the two phenotypes. Thus, for each transcriptomic dataset, we quantified the J-metric for the pairwise correlation matrix based on 14 NOR-specific and 12 MES-specific genes and calculated the correlation coefficient between the single-sample gene set enrichment analysis (ssGSEA) scores for the 359-gene NOR signature vs. those for 485-gene MES signature. We observed across datasets that the value of J-metric was positive, and the correlation coefficient between NOR and MES signatures was negative. We observed that the J-metric and correlation coefficients between the larger NOR/MES gene lists are negatively correlated with each other (r = - 0.33, p < 0.05, **Figure 3F**). This trend indicates that both the sets of genes can consistently capture the antagonism between the two phenotypes with a high degree of consistency, thereby suggesting that the ‘team-like’ behavior can be scalable. Put together, these results indicate that two distinct phenotypes can be observed across NB cell lines and primary tumor samples, along with the prevalence of two “teams” of genes driving each phenotype.

### ‘Teams-like’ behavior is exclusive to the set of NOR/MES specific genes

Our previous analysis revealed that the “teams” structure was largely unique to the underlying biological network. When compared to an ensemble of random networks of similar size and density, and same in-degree and out-degree of each node in the network, the biological network usually had a much higher team strength, thus highlighting that the “teams” topology is a salient feature of networks controlling cancer cell plasticity (Chauhan et al., 2021; Hari et al., 2022; Pillai and Jolly, 2021). Thus, we next investigated how exclusive the “teams-like” behavior is to the 14 NOR-specific and 12 MES-specific data sets.

As a control, we generated 1000 random ensembles of 26 genes from a set of housekeeping genes (**Table S2**) and performed PCA for the expression value matrix for those 26 genes in each dataset. We noticed that across datasets, the variance explained by PC1 was much higher for the ‘wild-type’ 26 gene set (14 NOR-specific and 12 MES-specific) than that explained by PC1 corresponding to the 1000 random combinations of 26 housekeeping genes (**Figure 4A, i-iii, S2**). As another control case, we generated 1000 random combinations of 26 genes with 14 of them randomly chosen from the 369 NOR-specific gene set and 12 of them randomly chosen from the 485 MES-specific gene set and quantified the variance along PC1 for all of them. On average, the ensemble of these 1000 combinations accounted for a higher PC1 variance than the ensemble of 1000 combinations comprising housekeeping genes. This trend suggests that the ‘teams-like’ behavior is exclusive to the specific sets of genes associated with a NOR and MES phenotype.

**Figure 4.**
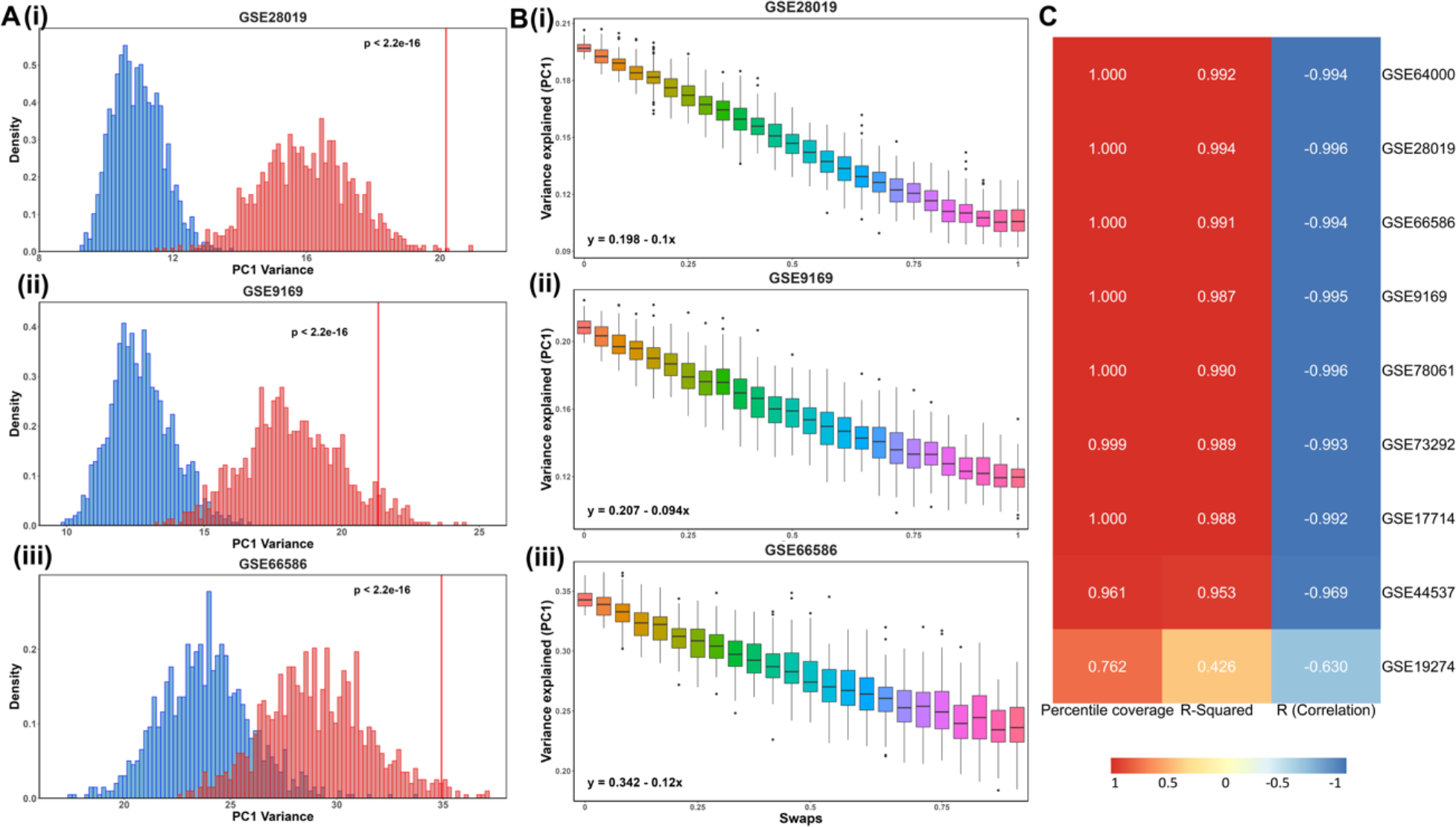
‘Team’-like behavior is exclusive to the set of NOR/MES team genes. **A)** i) Histogram (in red) showing the percentage of variance explained by PC1 for 1000 unique combinations of randomly chosen housekeeping genes for GSE28019. The vertical red line depicts the variance explained by the original NOR-MES 26 gene list. Histogram (in blue) showing the same percentage of variance explained by PC1 for 1000 unique combinations based on 14 genes chosen randomly from 369 NOR-specific signature, and 12 genes chosen randomly from 485 MES-specific signature. ii) and iii) are same as i) but for GSE9169 and GSE66586 respectively. **B)** i) Boxplots illustrating distribution of the fraction of variance explained by PC1 (y-axis) vs. number of genes from NOR/MES gene list swapped with housekeeping genes for (x-axis) for GSE28019. The equations for a linear fit of mean values for each swap are also shown. ii) and iii) same as i) but for GSE9169 and GSE66586 respectively. **C)** Heatmap depicting the percentile of variance explained by PC1 by original NOR-MES list in distribution of PC1 variance for random combinations of housekeeping genes; R-squared value (goodness of fit) and Pearson correlation coefficient for linear fit on mean values of variance explained by PC1 (Panels A,B).

To better understand this trend, we began with replacing one gene at a time in the 26-gene set (14 NOR-specific and 12 MES-specific) with a housekeeping gene. Given many possible combinations depending on which of the 26 genes is (are) being replaced and with which housekeeping gene(s), we tried 100 combinations for each scenario of a fixed number of swaps and plotted the distribution of PC1 variance corresponding to them. We noticed a linear decrease in the average PC1 variance for a given number of swaps, thus, the higher the no. of genes swapped from the 26-gene set, the lower the PC1 variance explained (**Figure 4B, S2**). This analysis endorses the low dimensionality of the NOR-MES phenotypic landscape and validated the presence of strong “teams” of players exclusive to the genes considered in the analysis. These trends break down once random housekeeping genes that are not connected with NOR-MES axis are introduced.

The trends of 1) a higher PC1 variance of the 26-gene set than a corresponding set of 26 housekeeping genes, and 2) a negative correlation between the number of genes swapped and the PC1 variance were both consistently observed across all nine datasets (**Figure 4C**) investigated (**Figure 1**).

### Experimental perturbations cause phenotypic alterations along the NOR-MES axis

To further substantiate the antagonistic enrichment of the two phenotypes in samples at a bulk level, we obtained ssGSEA scores of each sample across multiple datasets (denoted by GSE IDs) and compared scores of the samples with perturbations to that of the control/wildtype (**Figure 5**).

**Figure 5.**
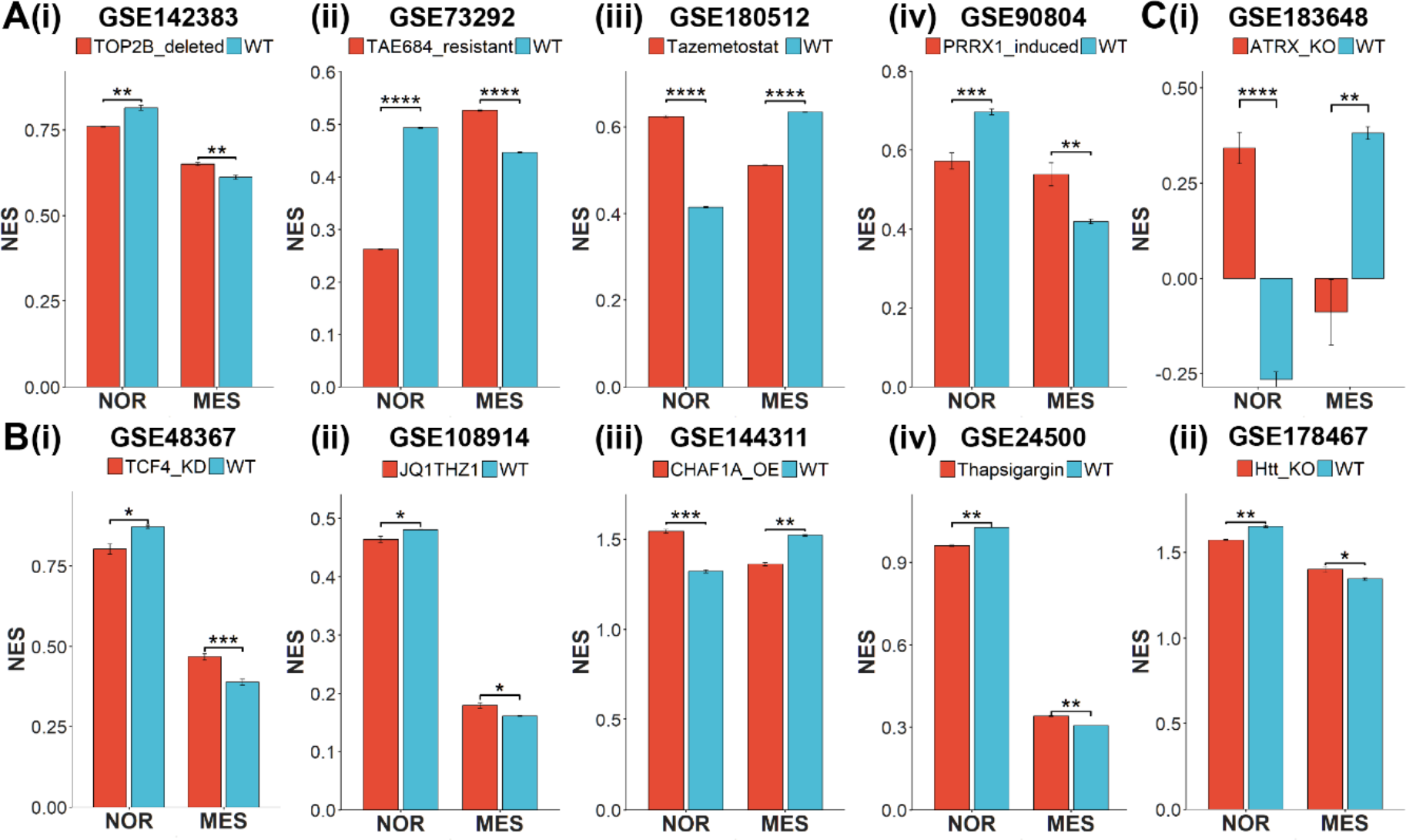
Experimental perturbations reveal changes in noradrenergic and mesenchymal programs in bulk RNA-seq data. Barplots depicting experimentally observed significant changes in ssGSEA scores of NOR and MES gene signatures in control samples vs. samples with different perturbations in bulk RNA-seq datasets denoted by respective GSE IDs. Error bars represent standard errors in mean values of the replicates and statistically significant differences are indicated by asterisks (*, **, ***, **** for p<0.05, <0.01, <0.001 and <0.0001 respectively) for Bonferroni-adjusted p-values.

Topoisomerase-2B (TOP2B) was shown to be essential for SH-SY5Y cells to maintain an adrenergic neural-like (NOR) phenotype, and its deletion upregulated mesenchymal (MES) markers (Khazeem et al., 2022). Consistent with this observation, RNA-seq data for TOP2B-null samples, relative to control, showed a significant reduction in enrichment of a 369 NOR-based gene list and a concomitant enrichment of a 485 MES-based gene list (**Figure 5A, i**) (GSE142383). Further, ALK inhibitor (TAE684)-resistant SH-SY5Y cells showed AXL activation and induction of EMT (Debruyne et al., 2016). Similar behavior was reflected in transcriptomic data where resistant SH-SY5Y cells were enriched in the MES gene set, and downregulated in the NOR gene set, when compared to the parental cell-line (**Figure 5A, ii**) (GSE73292). Next, SK-N-AS cells, upon treatment with the EZH2 inhibitor, tazemetostat, exhibit an increase in NOR scores and reduced MES scores (GSE180512 (**Figure 5A, iii**). PRRX1A is a MES-specific transcription factor whose overexpression in adrenergic-type SK-N-BE(2C) cells could reprogram them to a mesenchymal state (GSE90804) (van Groningen et al., 2019, 2017). This role of PRRX1A was recapitulated in ssGSEA scores of NOR and MES gene sets (**Figure 5A, iv**). Along similar lines, knockdown of TCF4, a crucial neurodevelopmental TF, in SH-SY5Y cells led to increased expression of many EMT regulators such as SNAI2, ZEB2 and other TGF-β targets (Forrest et al., 2013). These experimental observations are reinforced in transcriptomic analysis of enrichment of NOR and MES specific gene lists (**Figure 5B, i**) (GSE48367). Similar results were observed for BE2C cells treated with both JQ1 (a BET bromodomain inhibitor) and THZ1 (a CDK7 inhibitor) that decreased the expression levels of relevant transcription factors, including HAND2, PHOX2B, TBX2, ISL1, MYCN and GATA3 (Durbin et al., 2018). These six factors are a part of the 14 NOR specific gene set, thus their downregulation via JQ1 and THZ1, as expected, led to a reduced NOR signature and an increased MES signature (**Figure 5B, ii**) (GSE108914). Further, overexpression of chromatin assembly factor 1 subunit p150 (CHAF1A) – a known inhibitor of neuronal differentiation (Tao et al., 2021) – was able to significantly drive up the enrichment of the mesenchymal gene set in SHEP cells after 96 hours as compared to control (GSE144311) (**Figure 5B, iii**). We observed a similar trend with SH-SY5Y cells treated with thapsigargin, an endoplasmic reticulum Ca(2+)- ATPase inhibitor that is known to inhibit cell proliferation (Treiman et al., 1998). Given the association of NOR with a higher proliferative status than MES states (**Figure S3, Table S5**), we noticed a decrease in NOR scores and simultaneous increase in MES scores upon thapsigargin treatment (**Figure 5B, iv**) (GSE24500). Together, these results highlight how NOR and MES gene lists could capture the cell-state transitions driven by various cellular perturbations.

We next sought to identify additional factors associated with NMT/MNT. First, knockdown of ATRX in NGP (a neuroblastoma cell line) was found to enrich for NOR signature along with a decrease in MES signature (GSE183648) (**Figure 5C, i**), thus suggesting ATRX as a MES-specific factor. Second, knock out of Huntington’s gene (Htt) in SH-SY5Y NB cells upregulated the NOR gene signature with concurrent reduction in activity of MES signature, as compared to parental cells (**Figure 5C, ii**) (GSE178467) (Bensalel et al., 2021).

Overall, these results indicate that both known and novel experimental perturbations in NB cell lines can drive cell-state transitions along noradrenergic/mesenchymal spectrum in an antagonistic manner.

### Meta-analysis of transcriptomic datasets reveals association of NOR/MES phenotypes with metabolic programs and immune markers

Next, we conducted a meta-analysis of 75 bulk transcriptomic datasets (**Table S3**) using ssGSEA to 1) characterize the modifications in metabolic programs brought on by these cell state transitions and 2) analyze their associations with novel immunotherapeutic targets for NB patients.

First, we noticed that out of 42 datasets that showed significant correlation between NOR and MES signatures, 36 of them (85.7%) were negatively correlated, in support of our previous findings (**Figure 6A, left**). We next investigated the correlation of NOR and MES genes with a signature associated with PD-L1 activity in carcinomas (Dondero et al., 2016; Sahoo et al., 2021). Consistent with the results observed for carcinomas, out of 37 datasets in which NOR and PD-L1 scores were significantly correlated, 36 (97.29%) showed a negative correlation and 92.3% (36/39) datasets showed a positive correlation with the MES phenotype (**Figure 6A, middle, right**).

**Figure 6.**
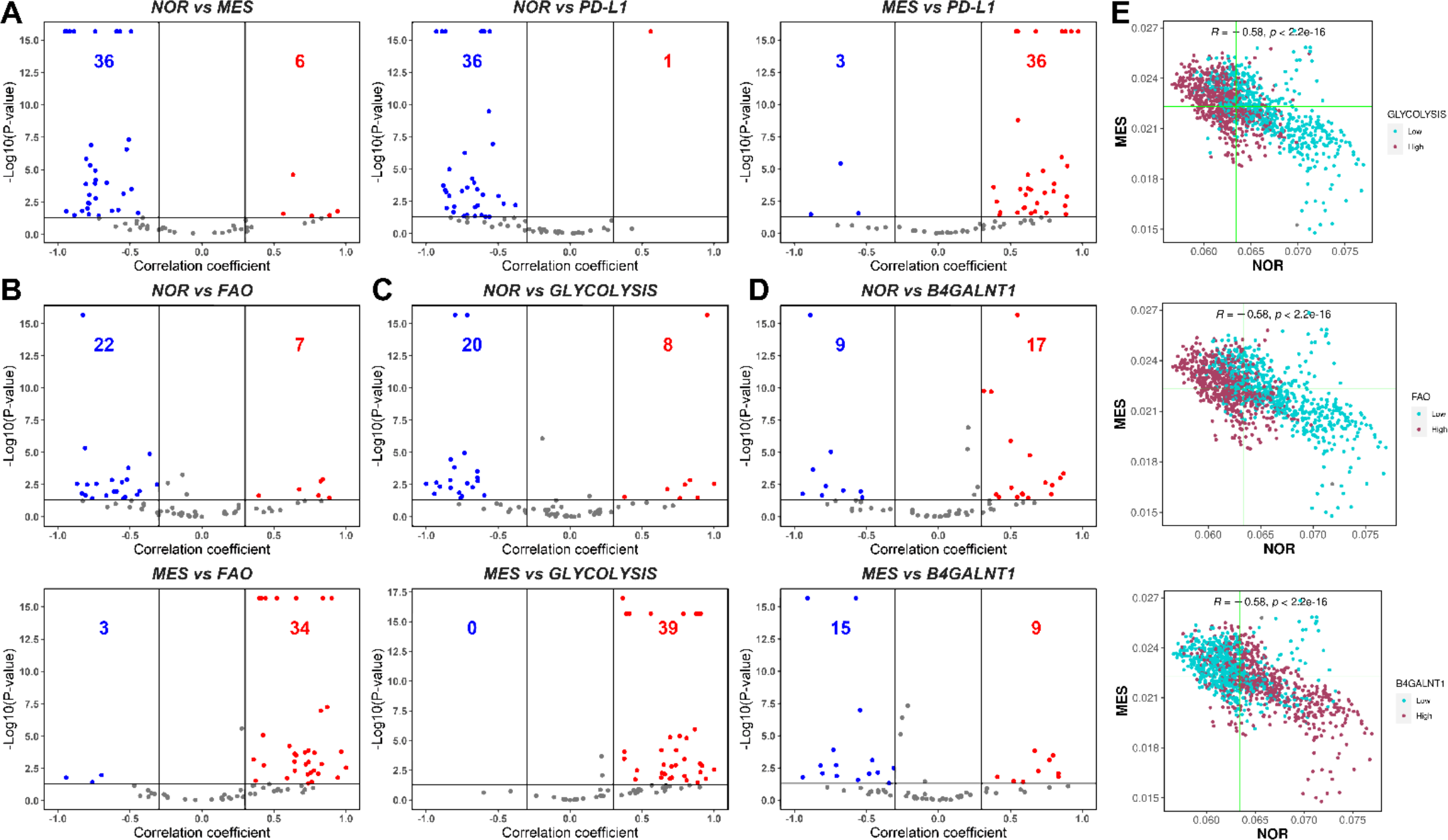
Modes of association between NOR and MES phenotypes with metabolic programs, PD-L1 and GD2 synthase gene (B4GALNT1). **A)** Volcano plots showing Spearman correlation coefficients (x-axis) and -log10(p-value) (y-axis) for NOR vs. MES (left), NOR vs. PD-L1 (middle) and MES vs. PD-L1 gene signature (right). Significant correlations (R > ± 0.3 and p < 0.05) are shown as red (positive) and blue (negative) datapoints. Same as **A)** but for **B)** NOR vs. FAO (top), MES vs. FAO (bottom), **C)** NOR vs. Glycolysis (top), MES vs. Glycolysis (bottom) and **D)** NOR vs. B4GALNT1 (top), MES vs. B4GALNT1 (bottom). **E)** Scatterplots of scRNA-seq data depicting NOR (x-axis) and MES scores (y-axis) of each cell for the control sample of GSE163429. The vertical and horizontal green lines are positioned at the median of NOR and MES scores, respectively. Maroon datapoints correspond to ‘high’, and blue datapoints represent ‘low’ Glycolysis (top), FAO scores (middle), imputed B4GALNT1 gene expression values (bottom). Threshold for ‘high’ and ‘low’ is set at the median value.

Additionally, we evaluated how these two phenotypes associated with two key metabolic processes — glycolysis and fatty acid oxidation (FAO) —that have been shown to be perturbed in NB cells (Slade et al., 2003; Zirath et al., 2013). In this context, we observed that in 75.9% datasets, the NOR phenotype is negatively linked with FAO, while the MES signature is strongly positively coupled with FAO (**Figure 6B**). Similarly, glycolysis scores are also negatively linked with NOR, but show a strong positive link with the MES phenotype. Further, activity of the NOR geneset correlated negatively with the Hallmark EMT signature while MES scores correlated positively, suggesting similarity between NMT and EMT at a transcriptomic level (**Figure S2A**). PD-L1 levels were found to be positively associated with both the hallmark EMT and partial EMT signatures, as well as glycolysis and FAO levels across datasets, reminiscent of observations made in carcinomas (Muralidharan et al., 2022; Sahoo et al., 2021) (**Figure S2**).

Finally, we also analyzed the links between NOR/MES scores with the cell surface marker protein—Disialoganglioside (GD2), which is abundantly expressed on cell surfaces of NB cells and is reported to be a potential target for immunotherapy in NB patients (Sait and Modak, 2017). However, recent observations have indicated that the mesenchymal state of NB cells can confer resistance to anti-GD2 therapy (Mabe et al., 2022). To characterize the association of GD2 with NMT, we used the gene expression values of *B4GALNT1* (GD2 synthase) – which showed a predominantly positive correlation with the NOR signature and a largely negative correlation with the MES signature, indicating a downregulation of GD2 in cells undergoing NMT. This observation may explain the observation of an enriched mesenchymal tumor population as a hallmark of anti-GD2 therapy resistance in NB patients.

To confirm our observations from bulk transcriptomic data, we also analyzed recently published scRNA-seq data of the NB cell line IMR-575 (Jansky et al., 2021). When projected onto the NOR-MES plane, cells showed a significant negative association (R = - 0.58, p < 10^-15^) between the two phenotypes (**Figure 6E**). These datapoints were then segregated into two groups – high and low – based on their ranks relative to the median AUCell enrichment score for a gene set. We observed that glycolysis^high^ cells were concentrated at the MES^high^NOR^low^ end of the plane, whereas glycolysis^low^ cells were predominantly present in the MES^low^NOR^high^ region (**Figure 6E, top**). FAO showed a similar trend —FAO^high^ cells were mostly clustered around the MES^high^NOR^low^ area and FAO^low^ cells were spread in MES^low^NOR^high^ portion of the 2D plane (**Figure 6E, middle**). Furthermore, B4GALNT1 expression was high for cells with MES^low^NOR^high^ whereas it was low in cells with MES^high^NOR^low^ phenotype (**Figure 6E, bottom**). These observations are largely concordant with our analysis of datasets at the bulk level and highlight the association of NOR/MES phenotypes with glycolysis, FAO and GD2 expression. These functional synergies might be leveraged in the future to design efficient combinatorial treatment methods that target multiple axes simultaneously to overcome therapy resistance.

## Discussion

NB is the most common form of extracranial cancer in infants (Brodeur, 2003). Its origin is attributed to improper differentiation of neural crest cells, and the degree of differentiation of these neural crest cells positively correlates with a favorable outcome for patients (Dong et al., 2020; Jansky et al., 2021; Kameneva et al., 2021; Rohrer, 2021). NB cells are heterogeneous in terms of biochemical and morphological traits, broadly classified into N-type (NOR), S-type (MES) and an intermediate (I-type) cell possessing traits of both N-and S-type cells (Biedler et al., 1973). These phenotypes have diverse transcriptomic and epigenetic profiles. While their interconversion has serious clinical implications, the dynamics of NMT/MNT is yet to be thoroughly studied (Gautier et al., 2021), unlike EMT/MET where systems-level computational and experimental approaches have mapped distinct cell-states, trajectories, and reversibility patterns (Tripathi et al., 2020).

To unravel the complexity and heterogeneity displayed by neuroblastoma, we collated a cohort of factors that are associated with the two major phenotypes (NOR/MES type) (van Groningen et al., 2017). Analyzing their expression levels in multiple bulk transcriptomic datasets pinpointed the presence of mutually exclusive ‘teams-like’ behavior among players that correspond to NOR and MES phenotypes. This behavior is reminiscent of our previous observations in other examples of cancer cell-state plasticity – melanoma, neuroendocrine differentiation and EMT (Chakraborty et al., 2021; Chauhan et al., 2021; Pillai and Jolly, 2021). PCA led to clear segregation of two clusters, which can be mapped to either a NOR or MES phenotype using GSEA. Importantly, the ‘teams-like’ behavior and consequent PC1 variance is specific to NOR and MES gene lists, and was not witnessed for housekeeping genes, indicating functional implications of this mutual antagonism. Such ‘teams’ behavior for epithelial (E) and mesenchymal (M) factors, but not for hybrid E/M ones, was proposed to underlie the difference in plasticity of these phenotypes (Hari et al., 2022). Similar experimental observations for I-type as reported for hybrid E/M phenotypes – higher plasticity, tumor-forming ability and malignancy, and correlation with worse survival as compared to N-type or S-type (Pasani et al., 2021; Pastushenko and Blanpain, 2019; Ross and Spengler, 2007; Walton et al., 2004) – and similar “teams-like” behavior seen in transcriptomic data for both EMT/MET and NMT/MNT highlight possible common fundamental design principles in regulatory networks and corresponding phenotypic space.

The NOR/MES antagonism was also distinctly observed in our meta-analysis of 75 bulk RNA-seq datasets, and in 10 datasets where we evaluated different experimental perturbations capable of inducing a phenotypic switch along the noradrenergic/mesenchymal axis. Functionally speaking, we observed a positive association of MES phenotype with both glycolysis and PD-L1 signatures, reminiscent of mesenchymal state traits seen in multiple carcinomas (Muralidharan et al., 2022). Expression levels of GD2, a cell surface marker reported to be a therapeutic target for NB patients (Sait and Modak, 2017), correlated negatively with MES, but positively with a NOR phenotype. These trends were also largely consistent at a single-cell level and suggest that combinatorial targeting of GD2 and PD-L1 may be a potent therapeutic strategy to overcome NB heterogeneity.

## Acknowledgements

This work was supported by Ramanujan Fellowship (SB/S2/RJN-049/2018) awarded to MKJ by Science and Engineering Research Board, Government of India.

## Author contributions

MKJ designed and supervised research, obtained funding. MS, SPN and SS performed research. All authors contributed to data analysis and writing of the manuscript.

## Supplementary figures

**Figure S1.**
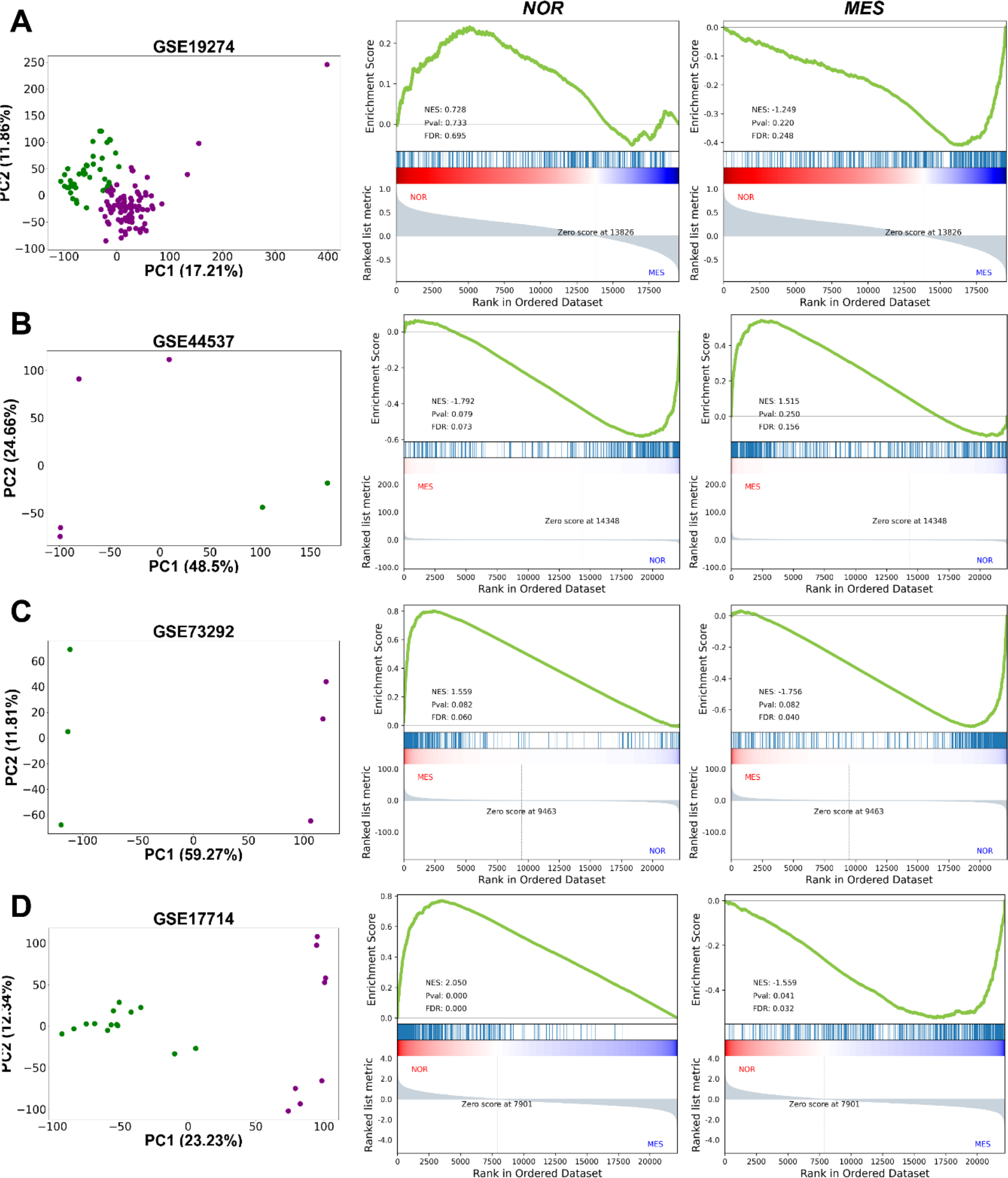
Two distinct classes of phenotypes—mesenchymal and adrenergic observed after dimensional reduction. **A)** PCA plot showing two distinct classes of samples exist in multiple datasets. K-means clustering for K = 2 yields two clusters (shown in green and violet color) along principal component 1 (PC-1). GSEA for undifferentiated mesenchymal (MES) geneset (middle) and differentiated adrenergic (NOR) geneset (right) confirms that these two clusters correspond to the respective phenotypes for **A)** GSE19274 (n = 138), **B)** GSE44537 (n = 6), **C)** GSE73292 (n = 6), **D)** GSE17714 (n = 22). ‘n’ stands for number of samples in each dataset. Percent variance explained by each principal component has been indicated along the respective axes.

**Figure S2.**
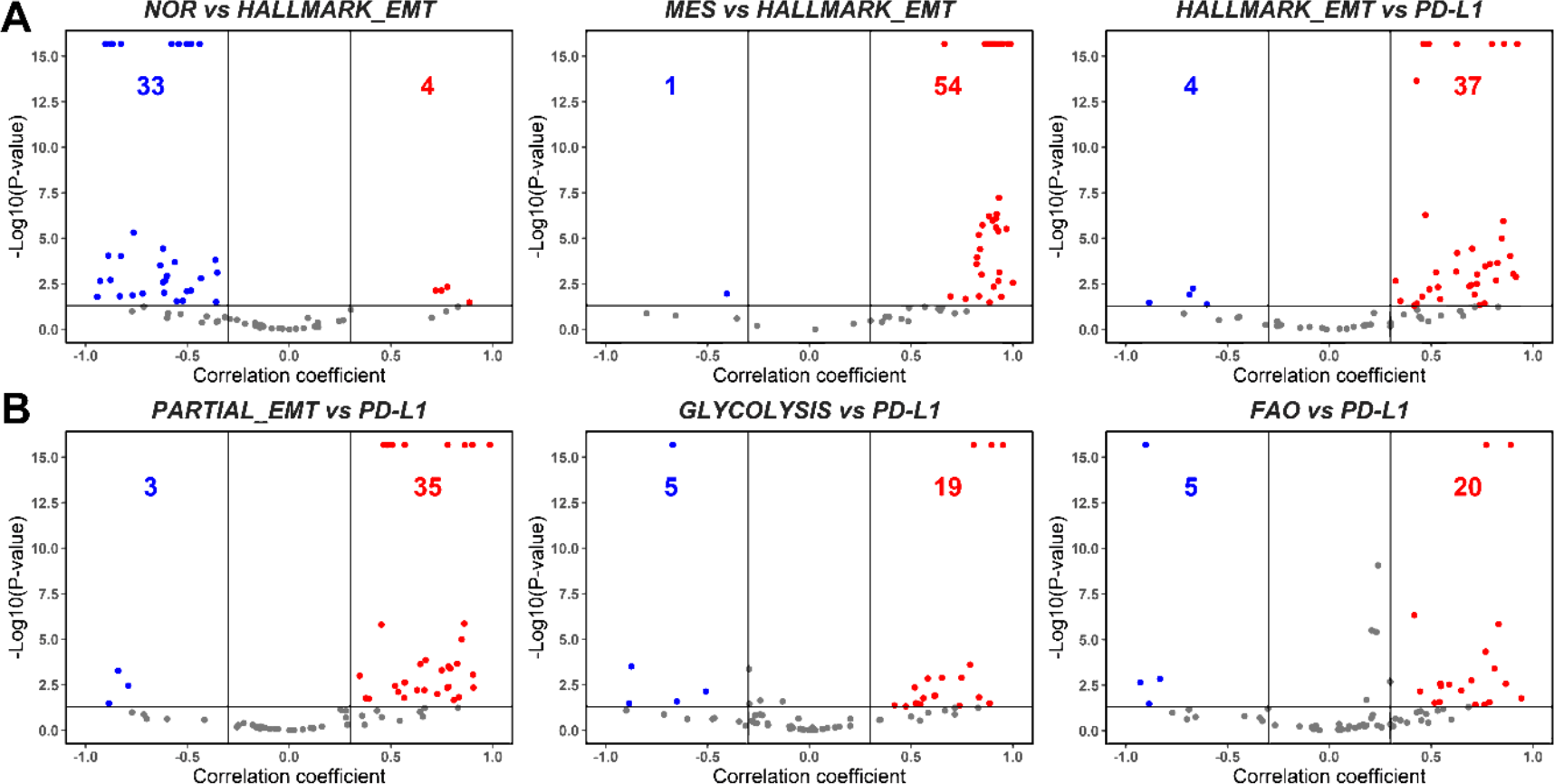
Modes of association between NOR/MES phenotypes with EMT, metabolism and PD-L1 in bulk transcriptomic data. **A)** Volcano plots showing spearman correlation coefficients (x-axis) and -log10(p-value) (y-axis) for NOR vs. Hallmark EMT (left), MES vs. Hallmark EMT (middle) and Hallmark EMT vs. PD-L1 gene signature (right). Significant correlations (R > ± 0.3 and p < 0.05) are shown as red (positive) and blue (negative) datapoints. Same as A) but for **B)** Partial EMT vs. PD-L1 (left), Glycolysis vs. PD-L1 (middle), and FAO vs. PD-L1 (right) scores.

**Figure S3.**
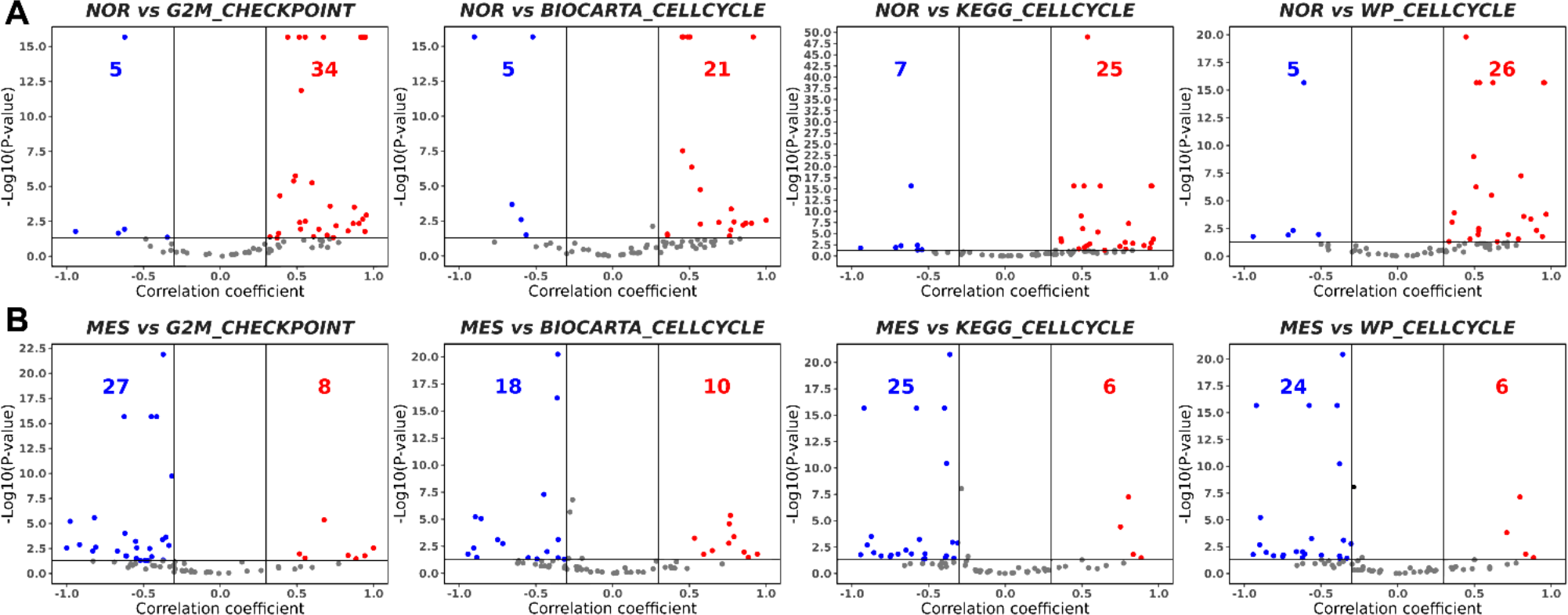
Modes of association between NOR/MES phenotypes with cell cycle genesets. **A)** Volcano plots showing spearman correlation coefficients (x-axis) and -log10(p-value) (y-axis) for NOR vs. Hallmark G2M checkpoint (left), NOR vs. Biocarta cell cycle geneset (middle-left), NOR vs. KEGG cell cycle gene signature (middle-right) and NOR vs. WP cell cycle signature (right). Significant correlations (R > ± 0.3 and p < 0.05) are shown as red (positive) and blue (negative) datapoints. Same as A) but for **B)** MES vs. Hallmark G2M checkpoint (left), NOR vs. Biocarta cell cycle (middle-left), NOR vs. KEGG cell cycle (middle-right) and NOR vs. WP cell cycle (right) scores.

**Figure S4.**
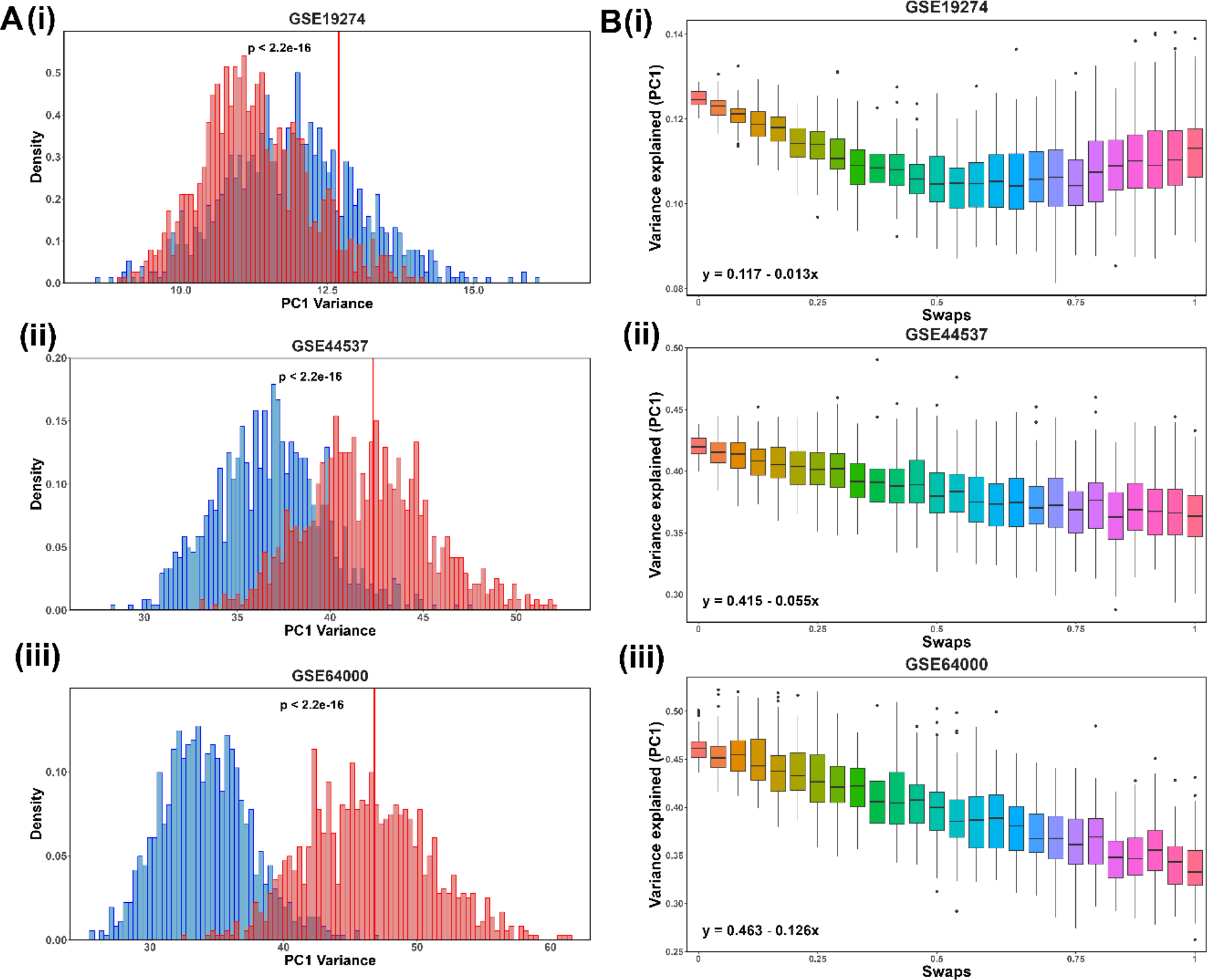
‘Team’-like behavior is exclusive to the set of NOR/MES team genes. **A)** i) Histogram (in red) showing the percentage of variance explained by PC1 for 1000 unique combinations of randomly chosen housekeeping genes for GSE19274. The vertical red line depicts the variance explained by the original NOR-MES 26 gene list. Histogram (in blue) showing the same percentage of variance explained by PC1 for 1000 unique combinations based on 14 genes chosen randomly from 369 NOR-specific signature, and 12 genes chosen randomly from 485 MES-specific signature. ii) and iii) are same as i) but for GSE44537 and GSE64000 respectively. **B)** i) Boxplots illustrating distribution of the fraction of variance explained by PC1 (y-axis) vs. number of genes from NOR/MES gene list swapped with housekeeping genes for (x-axis) for GSE28019. The equations for a linear fit of mean values for each swap are also shown. ii) and iii) same as i) but for GSE44537 and GSE64000 respectively.

## References

Aibar S, González-Blas CB, Moerman T, Huynh-Thu VA, Imrichova H, Hulselmans G, Rambow F, Marine J-C, Geurts P, Aerts J, van den Oord J, Atak ZK, Wouters J, Aerts S. 2017. SCENIC: single-cell regulatory network inference and clustering. Nat Methods 14:1083–1086. doi:10.1038/nmeth.4463

Bensalel J, Xu H, Lu ML, Capobianco E, Wei J. 2021. RNA-seq analysis reveals significant transcriptome changes in huntingtin-null human neuroblastoma cells. BMC Med Genomics 14:176. doi:10.1186/S12920-021-01022-W

Bhatia S, Monkman J, Blick T, Pinto C, Waltham A, Nagaraj SH, Thompson EW. 2019. Interrogation of Phenotypic Plasticity between Epithelial and Mesenchymal States in Breast Cancer. J Clin Med 8:893. doi:10.3390/jcm8060893

Biedler JL, Helson L, Spengler BA. 1973. Morphology and growth, tumorigenicity, and cytogenetics of human neuroblastoma cells in continuous culture. Cancer Res 33:2643–52.

Boeva V, Louis-Brennetot C, Peltier A, Durand S, Pierre-Eugène C, Raynal V, Etchevers HC, Thomas S, Lermine A, Daudigeos-Dubus E, Geoerger B, Orth MF, Grünewald TGP, Diaz E, Ducos B, Surdez D, Carcaboso AM, Medvedeva I, Deller T, Combaret V, Lapouble E, Pierron G, Grossetête-Lalami S, Baulande S, Schleiermacher G, Barillot E, Rohrer H, Delattre O, Janoueix-Lerosey I. 2017. Heterogeneity of neuroblastoma cell identity defined by transcriptional circuitries. Nature Genetics 2017 49:9 49:1408–1413. doi:10.1038/ng.3921

Brodeur GM. 2003. Neuroblastoma: biological insights into a clinical enigma. Nat Rev Cancer 3:203–216. doi:10.1038/nrc1014

Chakraborty P, Chen EL, McMullen I, Armstrong AJ, Jolly MK, Somarelli JA. 2021. Analysis of immune subtypes across the epithelial-mesenchymal plasticity spectrum. Comput Struct Biotechnol J 19:3842–3851. doi:10.1016/j.csbj.2021.06.023

Chauhan L, Ram U, Hari K, Jolly MK. 2021. Topological signatures in regulatory network enable phenotypic heterogeneity in small cell lung cancer. Elife 10:e64522. doi:10.7554/eLife.64522

Cole KA, Huggins J, Laquaglia M, Hulderman CE, Russell MR, Bosse K, Diskin SJ, Attiyeh EF, Sennett R, Norris G, Laudenslager M, Wood AC, Mayes PA, Jagannathan J, Winter C, Mosse YP, Maris JM. 2011. RNAi screen of the protein kinome identifies checkpoint kinase 1 (CHK1) as a therapeutic target in neuroblastoma. Proc Natl Acad Sci U S A 108:3336–3341. doi:10.1073/PNAS.1012351108/-/DCSUPPLEMENTAL/PNAS.201012351SI.PDF

Debruyne DN, Bhatnagar N, Sharma B, Luther W, Moore NF, Cheung NK, Gray NS, George RE. 2016. ALK inhibitor resistance in ALK(F1174L)-driven neuroblastoma is associated with AXL activation and induction of EMT. Oncogene 35:3681–3691. doi:10.1038/ONC.2015.434

Dong R, Yang R, Zhan Y, Lai H-D, Ye C-J, Yao X-Y, Luo W-Q, Cheng X-M, Miao J-J, Wang J-F, Liu B-H, Liu X-Q, Xie L-L, Li Y, Zhang M, Chen L, Song W-C, Qian W, Gao W-Q, Tang Y-H, Shen C-Y, Jiang W, Chen G, Yao W, Dong K-R, Xiao X-M, Zheng S, Li K, Wang J. 2020. Single-Cell Characterization of Malignant Phenotypes and Developmental Trajectories of Adrenal Neuroblastoma. Cancer Cell 38:716-733.e6. doi:10.1016/j.ccell.2020.08.014

Durbin AD, Zimmerman MW, Dharia N V., Abraham BJ, Iniguez AB, Weichert-Leahey N, He S, Krill-Burger JM, Root DE, Vazquez F, Tsherniak A, Hahn WC, Golub TR, Young RA, Look AT, Stegmaier K. 2018. Selective gene dependencies in MYCN-amplified neuroblastoma include the core transcriptional regulatory circuitry. Nat Genet 50:1240. doi:10.1038/S41588-018-0191-Z

Eisenberg E, Levanon EY. 2013. Human housekeeping genes, revisited. Trends Genet 29:569–74. doi:10.1016/j.tig.2013.05.010

Fardin P, Barla A, Mosci S, Rosasco L, Verri A, Versteeg R, Caron HN, Molenaar JJ, Øra I, Eva A, Puppo M, Varesio L. 2010. A biology-driven approach identifies the hypoxia gene signature as a predictor of the outcome of neuroblastoma patients. Mol Cancer 9. doi:10.1186/1476-4598-9-185

Forrest MP, Waite AJ, Martin-Rendon E, Blake DJ. 2013. Knockdown of human TCF4 affects multiple signaling pathways involved in cell survival, epithelial to mesenchymal transition and neuronal differentiation. PLoS One 8. doi:10.1371/JOURNAL.PONE.0073169

Gautier M, Thirant C, Delattre O, Janoueix-Lerosey I. 2021. Plasticity in neuroblastoma cell identity defines a noradrenergic-to-mesenchymal transition (Nmt). Cancers (Basel) 13. doi:10.3390/CANCERS13122904/S1

Greengard EG. 2018. Molecularly Targeted Therapy for Neuroblastoma. Children 5:142. doi:10.3390/CHILDREN5100142

Gu L, Chu P, Lingeman R, McDaniel H, Kechichian S, Hickey RJ, Liu Z, Yuan YC, Sandoval JA, Fields GB, Malkas LH. 2015. The Mechanism by Which MYCN Amplification Confers an Enhanced Sensitivity to a PCNA-Derived Cell Permeable Peptide in Neuroblastoma Cells. EBioMedicine 2:1923–1931. doi:10.1016/J.EBIOM.2015.11.016

Hari K, Harlapur P, Saxena A, Girish A, Levine H, Jolly MK. 2023. Low dimensionality of phenotypic space as an emergent property of coordinated teams in biological regulatory networks. bioRxiv 2023.02.03.526930. doi:10.1101/2023.02.03.526930

Hari K, Ullanat V, Balasubramanian A, Gopalan A, Jolly MK. 2022. Landscape of epithelial mesenchymal plasticity as an emergent property of coordinated teams in regulatory networks. Elife 11:e76535. doi:10.7554/elife.76535

Ikegaki N, Shimada H, Fox AM, Regan PL, Jacobs JR, Hicks SL, Rappaport EF, Tang XX. 2013. Transient treatment with epigenetic modifiers yields stable neuroblastoma stem cells resembling aggressive large-cell neuroblastomas. Proc Natl Acad Sci U S A 110:6097–6102. doi:10.1073/PNAS.1118262110/-/DCSUPPLEMENTAL/PNAS.201118262SI.PDF

Jansky S, Sharma AK, Körber V, Quintero A, Toprak UH, Wecht EM, Gartlgruber M, Greco A, Chomsky E, Grünewald TGP, Henrich K-O, Tanay A, Herrmann C, Höfer T, Westermann F. 2021. Single-cell transcriptomic analyses provide insights into the developmental origins of neuroblastoma. Nat Genet 53:683–693. doi:10.1038/s41588-021-00806-1

Johnsen JI, Dyberg C, Wickström M. 2019. Neuroblastoma—A neural crest derived embryonal malignancy. Front Mol Neurosci 12:433925. doi:10.3389/FNMOL.2019.00009/BIBTEX

Kahlert UD, Joseph J V., Kruyt FAE. 2017. EMT- and MET-related processes in nonepithelial tumors: importance for disease progression, prognosis, and therapeutic opportunities. Mol Oncol 11:860–877. doi:10.1002/1878-0261.12085

Kameneva P, Artemov A V., Kastriti ME, Faure L, Olsen TK, Otte J, Erickson A, Semsch B, Andersson ER, Ratz M, Frisén J, Tischler AS, de Krijger RR, Bouderlique T, Akkuratova N, Vorontsova M, Gusev O, Fried K, Sundström E, Mei S, Kogner P, Baryawno N, Kharchenko P V., Adameyko I. 2021. Single-cell transcriptomics of human embryos identifies multiple sympathoblast lineages with potential implications for neuroblastoma origin. Nat Genet 53:694–706. doi:10.1038/s41588-021-00818-x

Khazeem MM, Casement JW, Schlossmacher G, Kenneth NS, Sumbung NK, Chan JYT, McGow JF, Cowell IG, Austin CA. 2022. TOP2B Is Required to Maintain the Adrenergic Neural Phenotype and for ATRA-Induced Differentiation of SH-SY5Y Neuroblastoma Cells. Mol Neurobiol 59:5987–6008. doi:10.1007/s12035-022-02949-6

Lecca MC, Jonker MA, Kulsoom Abdul U, Uçuçüçükosmanoglu AK, Van Wieringen W, Westerman BA. 2018. Adrenergic to mesenchymal fate switching of neuroblastoma occurs spontaneously in vivo resulting in differential tumorigenic potential. Journal of Molecular and Clinical Medicine 2018, 1(4), 219–226 1:219–226. doi:10.31083/J.JMCM.2018.04.4221

Liberzon A, Subramanian A, Pinchback R, Thorvaldsdóttir H, Tamayo P, Mesirov JP. 2011. Molecular signatures database (MSigDB) 3.0. Bioinformatics 27:1739–1740. doi:10.1093/bioinformatics/btr260

Lourenco AR, Ban Y, Crowley MJ, Lee SB, Ramchandani D, Du W, Elemento O, George JT, Jolly MK, Levine H, Sheng J, Wong ST, Altorki NK, Gao D. 2020. Differential contributions of pre-And post-EMT tumor cells in breast cancer metastasis. Cancer Res 80:163–169. doi:10.1158/0008-5472.CAN-19-1427/653773/AM/DIFFERENTIAL-CONTRIBUTIONS-OF-PRE-AND-POST-EMT

Mabe NW, Huang M, Dalton GN, Alexe G, Schaefer DA, Geraghty AC, Robichaud AL, Conway AS, Khalid D, Mader MM, Belk JA, Ross KN, Sheffer M, Linde MH, Ly N, Yao W, Rotiroti MC, Smith BAH, Wernig M, Bertozzi CR, Monje M, Mitsiades CS, Majeti R, Satpathy AT, Stegmaier K, Majzner RG. 2022. Transition to a mesenchymal state in neuroblastoma confers resistance to anti-GD2 antibody via reduced expression of ST8SIA1. Nat Cancer 3:976–993. doi:10.1038/s43018-022-00405-x

Middelbeek J, Visser D, Henneman L, Kamermans A, Kuipers AJ, Hoogerbrugge PM, Jalink K, van Leeuwen FN. 2015. TRPM7 maintains progenitor-like features of neuroblastoma cells: implications for metastasis formation. Oncotarget 6:8760–8776. doi:10.18632/ONCOTARGET.3315

Muralidharan S, Sehgal M, Soundharya R, Mandal S, Majumdar SS, Yeshwanth M, Saha A, Jolly MK. 2022. PD-L1 Activity Is Associated with Partial EMT and Metabolic Reprogramming in Carcinomas. Current Oncology 2022, Vol 29, Pages 8285-8301 29:8285–8301. doi:10.3390/CURRONCOL29110654

Nishida Y, Adati N, Ozawa R, Maeda A, Sakaki Y, Takeda T. 2008. Identification and classification of genes regulated by phosphatidylinositol 3-kinase- and TRKB-mediated signalling pathways during neuronal differentiation in two subtypes of the human neuroblastoma cell line SH-SY5Y. BMC Res Notes 1. doi:10.1186/1756-0500-1-95

Park JR, Eggert A, Caron H. 2010. Neuroblastoma: Biology, Prognosis, and Treatment. Hematol Oncol Clin North Am 24:65–86. doi:10.1016/j.hoc.2009.11.011

Pasani S, Sahoo S, Jolly MK. 2021. Hybrid E/M phenotype(s) and stemness: a mechanistic connection embedded in network topology. J Clin Med 10:60. doi:10.1101/2020.10.18.341271

Pastushenko I, Blanpain C. 2019. EMT Transition States during Tumor Progression and Metastasis. Trends Cell Biol 29:212–226. doi:10.1016/j.tcb.2018.12.001

Pillai M, Jolly MK. 2021. Systems-level network modeling deciphers the master regulators of phenotypic plasticity and heterogeneity in melanoma. iScience 24:103111. doi:10.1016/j.isci.2021.103111

Puram S V., Tirosh I, Parikh AS, Patel AP, Yizhak K, Gillespie S, Rodman C, Luo CL, Mroz EA, Emerick KS, Deschler DG, Varvares MA, Mylvaganam R, Rozenblatt-Rosen O, Rocco JW, Faquin WC, Lin DT, Regev A, Bernstein BE. 2017. Single-Cell Transcriptomic Analysis of Primary and Metastatic Tumor Ecosystems in Head and Neck Cancer. Cell 171:1611-1624.e24. doi:10.1016/j.cell.2017.10.044

Rohrer H. 2021. Linking human sympathoadrenal development and neuroblastoma. Nat Genet 53:593–594. doi:10.1038/s41588-021-00845-8

Ross RA, Spengler BA. 2007. Human neuroblastoma stem cells. Semin Cancer Biol 17:241–247. doi:10.1016/j.semcancer.2006.04.006

Sahoo S, Nayak SP, Hari K, Purkait P, Mandal S, Kishore A, Levine H, Jolly MK. 2021. Immunosuppressive traits of the hybrid epithelial/mesenchymal phenotype. Front Immunol 12:797261. doi:10.3389/fimmu.2021.797261

Sait S, Modak S. 2017. Anti-GD2 immunotherapy for neuroblastoma. Expert Rev Anticancer Ther 17:889–904. doi:10.1080/14737140.2017.1364995

Sharma A, Merritt E, Hu X, Cruz A, Jiang C, Sarkodie H, Zhou Z, Malhotra J, Riedlinger GM, D. S. 2019. Non-Genetic Intra-Tumor Heterogeneity Is a Major Predictor of Phenotypic Heterogeneity and Ongoing Evolutionary Dynamics in Lung Tumors. Cell Rep 29:2164–2174.

Shendy NAM, Zimmerman MW, Abraham BJ, Durbin AD. 2022. Intrinsic transcriptional heterogeneity in neuroblastoma guides mechanistic and therapeutic insights. Cell Rep Med 3:100632. doi:10.1016/J.XCRM.2022.100632

Simeonov KP, Byrns CN, Clark ML, Norgard RJ, Martin B, Stanger BZ, Shendure J, McKenna A, Lengner CJ. 2021. Single-cell lineage tracing of metastatic cancer reveals selection of hybrid EMT states. Cancer Cell 39:1150-1162.e9. doi:10.1016/j.ccell.2021.05.005

Slade RF, Hunt DA, Pochet MM, Venema VJ, Hennigar RA. 2003. Characterization and inhibition of fatty acid synthase in pediatric tumor cell lines. Anticancer Res 23:1235–43.

Su Y, Bintz M, Yang Y, Robert L, Ng AHC, Liud V, Ribas A, Heath JR, Wei W. 2019. Phenotypic heterogeneity and evolution of melanoma cells associated with targeted therapy resistance. PLoS Comput Biol 15:e1007034. doi:10.1371/journal.pcbi.1007034

Subramanian A, Tamayo P, Mootha VK, Mukherjee S, Ebert BL, Gillette MA, Paulovich A, Pomeroy SL, Golub TR, Lander ES, Mesirov JP. 2005a. Gene set enrichment analysis: A knowledge-based approach for interpreting genome-wide expression profiles. Proceedings of the National Academy of Sciences 102:15545 LP –15550. doi:10.1073/pnas.0506580102

Subramanian A, Tamayo P, Mootha VK, Mukherjee S, Ebert BL, Gillette MA, Paulovich A, Pomeroy SL, Golub TR, Lander ES, Mesirov JP. 2005b. Gene set enrichment analysis: A knowledge-based approach for interpreting genome-wide expression profiles. Proc Natl Acad Sci U S A 102:15545–15550. doi:10.1073/pnas.0506580102

Tao L, Moreno-Smith M, Ibarra-García-Padilla R, Milazzo G, Drolet NA, Hernandez BE, Oh YS, Patel I, Kim JJ, Zorman B, Patel T, Kamal AHM, Zhao Y, Hicks J, Vasudevan SA, Putluri N, Coarfa C, Sumazin P, Perini G, Parchem RJ, Uribe RA, Barbieri E. 2021. CHAF1A Blocks Neuronal Differentiation and Promotes Neuroblastoma Oncogenesis via Metabolic Reprogramming. Advanced Science 8:2005047. doi:10.1002/advs.202005047

Thirant C, Peltier A, Durand S, Kramdi A, Louis-Brennetot C, Pierre-Eugène C, Gautier M, Costa A, Grelier A, Zaïdi S, Gruel N, Jimenez I, Lapouble E, Pierron G, Sitbon D, Brisse HJ, Gauthier A, Fréneaux P, Grossetête S, Baudrin LG, Raynal V, Baulande S, Bellini A, Bhalshankar J, Carcaboso AM, Geoerger B, Rohrer H, Surdez D, Boeva V, Schleiermacher G, Delattre O, Janoueix-Lerosey I. 2023. Reversible transitions between noradrenergic and mesenchymal tumor identities define cell plasticity in neuroblastoma. Nature Communications 2023 14:1 14:1–18. doi:10.1038/s41467-023-38239-5

Treiman M, Caspersen C, Christensen SB. 1998. A tool coming of age: thapsigargin as an inhibitor of sarco-endoplasmic reticulum Ca2+-ATPases. Trends Pharmacol Sci 19:131–135. doi:10.1016/S0165-6147(98)01184-5

Tripathi S, Levine H, Jolly MK. 2020. The Physics of Cellular Decision-Making during Epithelial-Mesenchymal Transition. Annu Rev Biophys 49:1–18. doi:10.1146/annurev-biophys-121219-081557

van Dijk D, Sharma R, Nainys J, Yim K, Kathail P, Carr AJ, Burdziak C, Moon KR, Chaffer CL, Pattabiraman D, Bierie B, Mazutis L, Wolf G, Krishnaswamy S, Pe’er D. 2018. Recovering Gene Interactions from Single-Cell Data Using Data Diffusion. Cell 174:716-729.e27. doi:10.1016/j.cell.2018.05.061

van Groningen T, Akogul N, Westerhout EM, Chan A, Hasselt NE, Zwijnenburg DA, Broekmans M, Stroeken P, Haneveld F, Hooijer GKJ, Savci-Heijink CD, Lakeman A, Volckmann R, van Sluis P, Valentijn LJ, Koster J, Versteeg R, van Nes J. 2019. A NOTCH feed-forward loop drives reprogramming from adrenergic to mesenchymal state in neuroblastoma. Nat Commun 10. doi:10.1038/S41467-019-09470-W

Van Groningen T, Koster J, Valentijn LJ, Zwijnenburg DA, Akogul N, Hasselt NE, Broekmans M, Haneveld F, Nowakowska NE, Bras J, Van Noesel CJM, Jongejan A, Van Kampen AH, Koster L, Baas F, Van Dijk-Kerkhoven L, Huizer-Smit M, Lecca MC, Chan A, Lakeman A, Molenaar P, Volckmann R, Westerhout EM, Hamdi M, Van Sluis PG, Ebus ME, Molenaar JJ, Tytgat GA, Westerman BA, Van Nes J, Versteeg R. 2017. Neuroblastoma is composed of two super-enhancer-associated differentiation states. Nature Genetics 2017 49:8 49:1261–1266. doi:10.1038/ng.3899

van Groningen T, Koster J, Valentijn LJ, Zwijnenburg DA, Akogul N, Hasselt NE, Broekmans M, Haneveld F, Nowakowska NE, Bras J, van Noesel CJM, Jongejan A, van Kampen AH, Koster L, Baas F, van Dijk-Kerkhoven L, Huizer-Smit M, Lecca MC, Chan A, Lakeman A, Molenaar P, Volckmann R, Westerhout EM, Hamdi M, van Sluis PG, Ebus ME, Molenaar JJ, Tytgat GA, Westerman BA, van Nes J, Versteeg R. 2017. Neuroblastoma is composed of two super-enhancer-associated differentiation states. Nat Genet 49:1261–1266. doi:10.1038/ng.3899

Walton JD, Kattan DR, Thomas SK, Spengler BA, Guo HF, Biedler JL, Cheung NK V., Ross RA. 2004. Characteristics of Stem Cells from Human Neuroblastoma Cell Lines and in Tumors. Neoplasia 6:838. doi:10.1593/NEO.04310

Zage PE. 2018. Novel Therapies for Relapsed and Refractory Neuroblastoma. Children 5:148. doi:10.3390/CHILDREN5110148

Zeineldin M, Patel AG, Dyer MA. 2022. Neuroblastoma: When differentiation goes awry. Neuron 110:2916–2928. doi:10.1016/j.neuron.2022.07.012

Zirath H, Frenzel A, Oliynyk G, Segerström L, Westermark UK, Larsson K, Munksgaard Persson M, Hultenby K, Lehtiö J, Einvik C, Påhlman S, Kogner P, Jakobsson P-J, Arsenian Henriksson M. 2013. MYC inhibition induces metabolic changes leading to accumulation of lipid droplets in tumor cells. Proceedings of the National Academy of Sciences 110:10258–10263. doi:10.1073/pnas.1222404110

